# The BMP antagonist *Gremlin1* contributes to the development of cortical excitatory neurons, motor balance and fear responses

**DOI:** 10.1101/2020.07.24.219394

**Authors:** Mari Ichinose, Nobumi Suzuki, Tongtong Wang, Hiroki Kobayashi, Laura Vrbanac, Jia Q Ng, Josephine A Wright, Tamsin R M Lannagan, Krystyna A Gieniec, Martin Lewis, Ryota Ando, Atsushi Enomoto, Simon Koblar, Paul Thomas, Daniel L Worthley, Susan L Woods

## Abstract

Bone morphogenetic protein (BMP) signaling is required for early forebrain development and cortical formation. How the endogenous modulators of BMP signaling regulate the structural and functional maturation of the developing brain remains unclear. Here we show that expression of the BMP antagonist, *Grem1*, marks a neuroprogenitor that gives rise to layer V and VI glutamatergic neurons in the embryonic mouse brain. Lineage tracing of *Grem1*-expressing cells in the embryonic brain was examined by administration of tamoxifen to pregnant *Grem1creERT Rosa26LSLTdtomato* mice at 13.5 days post coitum (dpc), followed by collection of embryos later in gestation. In addition, at 14.5 dpc, bulk mRNA seq analysis of differentially expressed transcripts between FACS sorted *Grem1* positive and negative cells was performed. We also generated *Emx1-cre* mediated *Grem1* conditional knockout mice (*Emx1-Cre;Grem1^flox/flox^*) in which the *Grem1* gene was deleted specifically in the dorsal telencephalon. *Grem1^Emx1cKO^* animals had reduced cortical thickness, especially layers V and VI and impaired motor balance and fear sensitivity compared to littermate controls. This study has revealed new roles for Grem1 in the structural and functional maturation of the developing cortex.

**Summary statement:** The BMP antagonist, *Grem1*, marks neuroprogenitors that give rise to deep layer glutamatergic neurons in the embryonic mouse brain. *Grem1* conditional knockout mice display cortical and behavioural abnormalities.

## Introduction

During development of the nervous system, bone morphogenetic protein (BMP) signaling has important roles in the promotion of dorsal identity, and the regulation of cell proliferation and differentiation. Yet BMP function in the maturation of neurons in vivo is poorly understood. Misregulation of BMP signaling has been suggested to contribute to human neurodevelopmental conditions such as autism spectrum disorders (Kumar et al., 2019), however a detailed understanding of the exact role for BMP signaling in these disorders is lacking, in line with our imperfect knowledge of the regulation of the BMP pathway in normal brain development.

The BMP ligands (BMP 2, 4, 5, 6 and 7) bind to type I (BmprIA, BmprIB, Acvr1) and type II (BmprII, ActrIIA, and ActrIIB) receptors. BMP ligands are expressed broadly in the telencephalon during early gestation in mice and then become more tightly localized to the choroid plexus at 13.5 dpc (Furuta et al., 1997). BMPs have also been derived from the mesenchyme of the brain, such as meninges and endothelial cells (Imura et al., 2008, Choe and Pleasure, 2018). Both type I and II BMP receptors are expressed in the telencephalon during development but this is further restricted in adulthood to BmprII in the cortex and hippocampus and ActrIIA/IIB in the dentate gyrus (Söderström et al., 1996). Aside from this spatiotemporal regulation of receptors, the BMP signaling pathway is also regulated by a family of secreted extracellular antagonists, that directly bind to the BMP ligands to prevent interactions with BMP receptors both in development and disease (Ali and Brazil, 2014). Antagonists such as Gremlin1 (Grem1), Noggin (Nog) and Chordin (Chrd) have been shown to inhibit BMP action in a range of different cell types and developmental stage-specific contexts to provide exquisite spatiotemporal regulation of the pathway. The roles of Nog and Chrd have been partially elucidated: they are required for forebrain development (Bachiller et al., 2000), as well as to create a niche for adult hippocampal neurogenesis (Lim et al., 2000, Sun et al., 2007). The expression and function of Grem1 in the developing brain has not yet been determined. Grem1 is an extracellular secreted antagonist of BMP 2,4, and 7 that signals to intestinal stem cells in the gut (Worthley et al., 2015) and plays a crucial role in *Xenopus* dorsalisation (Hsu et al., 1998) and limb and kidney formation (Khokha et al., 2003). Likewise the role of Grem1 in the normal adult CNS is unmapped territory to date, outside of prior work in the pathogenic state of glioma (Yan et al., 2014, Guan et al., 2017, Fu et al., 2018).

BMP signaling has been implicated in the regulation of forebrain patterning during early embryogenesis, working together with other signaling pathways such as fibroblast growth factor (FGF), Wnt, and Notch. High levels of BMP activity suppresses anterior neural development, whereas abrogation of BMP signaling can promote neural specification (Wilson and Houart, 2004, Lamb et al., 1993, Bachiller et al., 2000). *In vitro* the addition of BMP 2 and 4 to mouse neural stem cell cultures represses cell proliferation (Mathieu et al., 2008). BMP ligands regulate neuronal differentiation as well as determination of glial cell fate, by promoting astrocyte differentiation at the expense of oligodendrocytic fates (Mathieu et al., 2008, Yun et al., 2004, Sun and Xu, 2010, Gomes et al., 2003).

Cortical neural stem cells (also known as radial glial cells, RGC) begin to divide asymmetrically to start producing neurons at around 11.5 dpc in mice, signaling the start of cortical development. RGC daughter cells detach from the ventricle and form the first neuronal layer of the preplate by 13.5 dpc (Martynoga et al., 2012). Subsequently born neurons migrate along RGCs and start to form the cortical plate, separating the preplate into Cajal-Retzius cells in the marginal zone and subplate neurons (Dwyer et al., 2016). The neocortex develops in an inside out manner, with deep layers emerging first and superficial layers last. These neurons differentiate into glutamatergic pyramidal neurons, whereas inhibitory interneurons are born in the subcortical ganglionic eminences. Deep-layer pyramidal neurons (DLPNs, layer V and VI) have both intratelenchephalic projections to superficial cortical layers and extracephalic projections to other brain regions (Baker et al., 2018). Multiple molecular mechanisms regulate this corticogenesis (Martynoga et al., 2012), however the role of BMP signaling in cortical layer formation and functional maturation of neurons has only been reported in limited studies so far. *In utero* electroporation of BMP7 to murine 14.5 dpc cortical ventricular cells impaired neuronal migration, suggesting BMP signaling regulates neuronal positioning and migration (Choe and Pleasure, 2018). BMPs also regulate dendritogenesis and neurite growth *in vitro* (Lee – Hoeflich et al., 2004, Matsuura et al., 2007). This is consistent with a recent study that suggests perturbation of BMP signaling by delivery of a dominant-negative version of BMP IB receptor affects migration, polarity and dendritogenesis of mouse cortical neurons *in vivo* (Saxena et al., 2018).

To further establish the role of BMP signaling in forebrain development and neuronal function, we focus here on the expression and function of the BMP antagonist Grem1 in the developing mouse brain. We first assess *Grem1* expression using transgenic *Grem1* reporter mice in the dorsal telencephalon and developing neocortex. Next, to investigate the role of *Grem1* in the developing brain we use transcriptomic analyses of sorted mouse *Grem1*-expressing cells and human single cell RNA sequence (scRNAseq) data, combined with mouse neural stem/progenitor cell (NSPC) culture *ex vivo*. Lastly, to examine the functional contribution of *Grem1* to cortical development, we conditionally delete *Grem1* in the dorsal telecephalon and undertake behavioural testing of mutant animals and littermate controls.

## Results

### *Grem1* expressing cells are located in the dorsal telencephalon and give rise to deep layer neocortical neurons

To assess *Grem1* expression in the embryonic mouse brain we utilized transgenic *Grem1creERT; Rosa26LSLTdtomato* reporter mice, in which tamoxifen treatment results in expression of TdTomato in cells where the *Grem1* enhancer and promoter sequences are active and the progeny of those cells (Worthley et al., 2015). Pregnant *Grem1creERT;Rosa26LSLTdtomato* mice were administered tamoxifen at 11.5 dpc and embryonic brains were collected 24 h later, but no TdTomato^+^ cells were observed, suggesting that *Grem1* is not yet expressed in the brain at this early time-point (**Suppl Fig.1A**). Next, we administered tamoxifen to pregnant dams at 13.5 dpc, and collected embryonic brains at 14.5, 17.5 and 20.5 dpc (**Fig.1A,B**). From this timepoint we observed cells expressing TdTomato 24 hours after tamoxifen administration located in the lower cortical plate and subplate of the dorsal telencephalon, with dendrites extending to the pia matter (**Fig.1B,G**). ISH confirmed that *Grem1* RNA was detected in almost all TdTomato^+^ cells at 14.5 dpc, confirming that the reporter line recapitulates endogenous *Grem1* expression. Six days later at 20.5 dpc, TdTomato^+^ cell somas had migrated toward the lateral ventricle with dendrites extending to the pia matter, becoming layer V pyramidal neurons (**Fig.1B,C**). Layer VI neurons were TdTomato^+^ at 17.5 dpc and expanded in number at 20.5 dpc (**Fig.1B,C**). ISH showed *Grem1* RNA was detected in most TdTomato^+^ layer V pyramidal neurons, but only 22% of layer VI neurons (**Fig.1C,D**). This suggests that *Grem1* is actively expressed in layer V neurons but not layer VI pyramidal neurons at 20.5 dpc. Cortical *Grem1* mRNA levels dramatically decrease after birth as determined by qRT-PCR analysis of total RNA isolated from mouse brain cortex at birth, p10 and 4w post-birth (**Fig.S1B**).

**Fig. 1.**
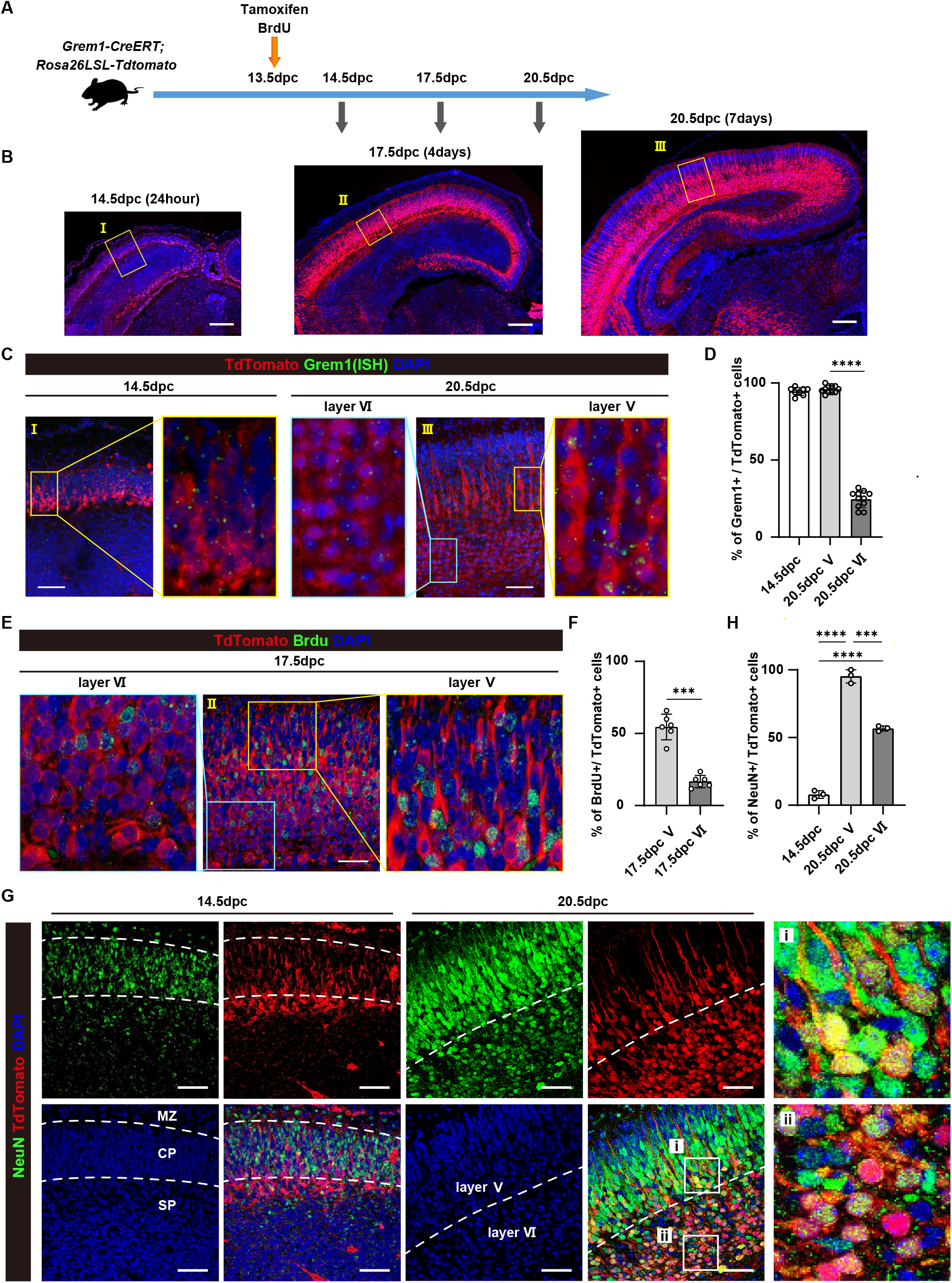
*Grem 1*-expressing cells give rise to cortical neurons in the developing mouse brain. (A) Schematic showing tamoxifen and BrdU administration to *Grem1creERT; R°sa26LSLTd”mat° (Grem1reporter)* mice. (B) Representative images of TdTomato^+^ (red) cells in the telencephalon at 44.5dpc (24h post-induction), 17.5dpc (3d post-induction), and 20.5dpc (6d post-induction) in *Grem1*-reporter mice treated with tamoxifen at 13.5 dpc, DAPI (blue). Scale bar = 200 μm. (C) Representative images of immunohistochemical staining of boxed region from (B) to visualize TdTomato^+^ cells (red) and *Grem1* mRNA by ISH (green) at 14.5dpc (I, 24h post-induction) and 20.5dpc (III, 6d post-induction-in *Grem1reporter* mice treated with tamoxifen at 13.5 dpc, DAPI (blue). The boxed areas were further magnified in adjacent panels. Scale bar = 50 μm. (D) Quantification of (C) showing the percentage of TdTomato^+^ cells that were also *Grem1* RNA^+^ in 4 HPFs of 2-3 biological replicates, t-test. (E) Representative images of immunofluorescence staining of (II) 17.5dpc telencephalon from *Grem1-reporter* (red) mice induced with tamoxifen at dpc13.5, BrdU (green), DAPI (blue). Scale bar = 50 μm. (F) Quantification of (E) showing the percentage of TdTomato^+^ cells that were also BrdU^+^ 2 HPFs of 3 biological replicates, t-test. (G) Representative images of immunofluorescence staining of 14.5 and 20.5 dpc neocortex from *Grem1reporter* (red) mice induced with tamoxifen at dpc13.5, NeuN (green), DAPI (blue). Layer V and VI boxed in (i) and (ii), respectively. Scale bar = 50 μm. MZ, marginal zone; CP, cortical plate; SP, subplate (H) Quantification of (G) showing the percentage of TdTomato^+^ cells that were also NeuN^+^ in 3 HPF from 3 biological replicates. One way ANOVA with Tukey’s multiple test. *p<0.05, **p<0.01, ***p<0.001, ****p<0.0001

To confirm that *Grem1* + cells are actively dividing in order to give rise to their traced progeny, we administered the thymidine analogue BrdU concomitantly with tamoxifen to pregnant *Grem1creERT; Rosa26LSLTdtonato* dams at 13.5 dpc (**Fig.1A**). BrdU is incorporated into newly synthesized DNA and so marks cells that are dividing. 55% of TdTomato^+^, layer V pyramidal neurons cells are labelled with BrdU at 17.5 dpc, demonstrating *Grem1* expressing cells at 13.5 dpc are progenitor cells undergoing mitosis (**Fig.1E,F**). Fewer TdTomato^+^ cells (17%) in layer VI were BrdU positive, suggesting lower cell proliferation over the same period (**Fig.1F**). We subsequently assessed whether TdTomato^+^ cells express a marker of post-mitotic mature neurons, NeuN, by immunohistochemistry. At 14.5 dpc most TdTomato^+^ cells were NeuN negative and were located in the subplate, and lower NeuN positive cortical plate (**Fig.1G**). In contrast, later in gestation at 20.5 dpc almost all TdTomato^+^ cells in layer V and approximately half of those in layer VI express NeuN (**Fig.1H**). This suggests that Grem1-lineage traced cells give rise to mature neurons in the telencephalon.

### Global transcriptomic analysis defines transcript modules enriched in *Grem1*-expressing cells

To further characterise *Grem1* expressing cells in the developing mouse brain, we undertook mRNAseq analysis on TdTomato^+^ and negative cells from the telencephalon. Pregnant *Grem1creERT; Rosa26LSLTdtomato* mice were administered tamoxifen at 13.5 dpc and embryonic brains were collected at 14.5 dpc. At this time-point almost all TdTomato^+^ cells express endogenous *Grem1* RNA (**Fig.1C**). Dissociated brain cells were flow sorted and live, TdTomato^+^ and TdTomato^-^ cells were collected for transcriptomic analysis. TdTomato^+^ cells accounted for 5.1 ± 2.1% of all live cells (n=5) (**Fig.2A**). Bulk mRNA sequencing of TdTomato^+^ and TdTomato^-^ populations and analysis of 1845 DEG between the two cell populations revealed that *Grem1* was significantly upregulated in TdTomato^+^ cells and BMP transcriptional target genes (Id1, Id3, Id4) were significantly downregulated (adjusted P value (FDR) ≤ 0.05 (**Fig.2B**). We next undertook a correlation analysis to identify modules of the 1845 DEGs that are coordinately regulated in TdTomato^+^ cells (**Fig.S2).** Shared membership of a module can suggest genes that together perform a particular function. The clustering tree depicts the topological distance between different modules, i.e. how similar or different the expression of transcripts within the module are from other modules and highlighted one module that contained 288 genes with particularly tightly correlated transcripts (light green module, **Fig.S2).** Network visualization of the correlation analysis showed that the majority of transcripts within this tightly correlated module are upregulated in TdTomato^+^ cells (magenta square, **Fig.2C)**. We investigated the known function of genes within this module using a hypogeometric test for gene set enrichment. Significantly enriched pathways (adjusted P value (FDR) ≤ 0.001) were related to neuronal differentiation and projection, calcium signaling (possibly associated with synapse functions) and axons (**Fig.2D**). The BMP target gene, *Id1*, was identified as one of the central hub genes in this cluster and its expression was significantly associated with other transcripts that have functions in neuronal maturation (such as Lrrtm3, Ryr3) (**Fig.2E**). This suggests that neuronal functions are upregulated in the *Grem1-*expressing TdTomato^+^ cells of the developing telencephalon, and that this biological function is linked, at least in part, to BMP signaling.

**Fig. 2.**
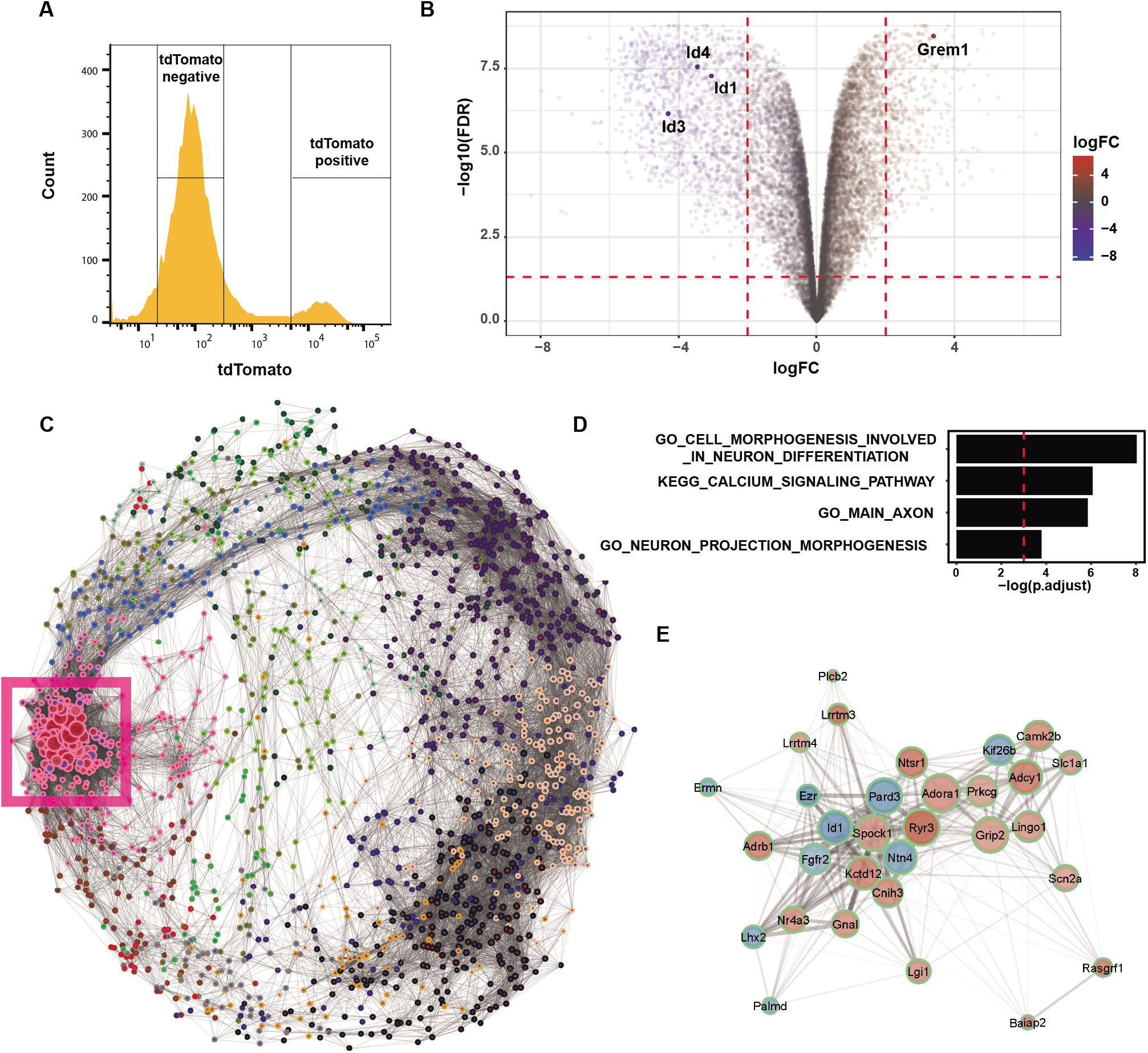
Grem1 expressing cells have neuron associated gene signatures. (A) TdTomato^+^ and TdTomato^-^ cells were isolated from 14.5 dpc brains of *Grem1-CreERT; R26-TdTomato* mice induced with tamoxifen at 13.5dpc. Representative FACS plot is shown, n=5 mice sorted for bulk RNAseq analysis. (B) Volcano plot to show differentially expressed genes (DEG) between TdTomato^+^.and TdTomato^-^ cells from (A). *Grem1* was significantly upregulated, whereas BMP target genes, *Id1,3,4* were significantly downregulated in TdTomato^+^ cells compared to TdTomato^-^ cells. Absolute value of log2 fold change ≤ 2.0, FDR<0.05 (C) Network visualization of correlated DEG modules in TdTomato^+^ cells. Each dot represents a gene, dot perimeter colour indicates module membership (see Fig S3), dot interior colour denotes upregulation (red) or downregulation (blue) of gene transcript in TdTomato^+^ cells compared to TdTomato^-^, dot size indicates the magnitude of gene expression correlation to neighboring genes, lines connecting dots represent topological distance. The module containing highly correlated genes that were predominantly upregulated in TdTomato^+^ cells is boxed in magenta. (D) Significantly enriched gene sets in magenta boxed module in (C). FDR<0.001 (E) Genes associated with Id1, a hub gene in the magenta boxed module in (C). Dot perimeter colour indicates module membership (see Fig S3), dot interior colour denotes upregulation (red) or downregulation (blue) of gene transcript in TdTomato^+^ cells compared to TdTomato).

### *Grem1/GREM1* is enriched in excitatory neuronal lineage cells during brain development

To further characterize the *Grem1-*expressing TdTomato^+^ cell population we undertook a candidate gene profiling approach using the bulk mRNAseq data. Gene expression patterns of neuronal markers from a series of differentiation stages are shown in **Fig.3A**. Immature neuronal markers, such as *Dcx, Ncam1, Neurod 2/6, Tbr1*, were significantly upregulated at the RNA level in TdTomato^+^ cells in comparison to the TdTomato^-^ cells at 14.5 dpc, while neural stem cell, radial glia, and intermediate progenitor transcripts were significantly under-represented (p<0.05). When RGCs (marked by co-expression of Sox2 and Pax6-generate neocortical neurons, the expression of specific transcription factors can be used to identify neurons within particular cortical layers(Martynoga et al., 2012). Our bulk RNAseq analysis showed that transcripts encoding the layer V marker, *Fezf2*, and VI markers, *Tbr1* / *Sox5*, were significantly upregulated in *Grem1*-expressing TdTomato^+^ cells, whereas the RGC markers, *Sox2* and *Pax6* were significantly downregulated (absolute value of log2 fold change≥2.0, FDR<0.05) and did not colocalize with Tdtomato at 14.5 dpc by immunohistochemistry (**Fig.S3**).

**Fig. 3.**
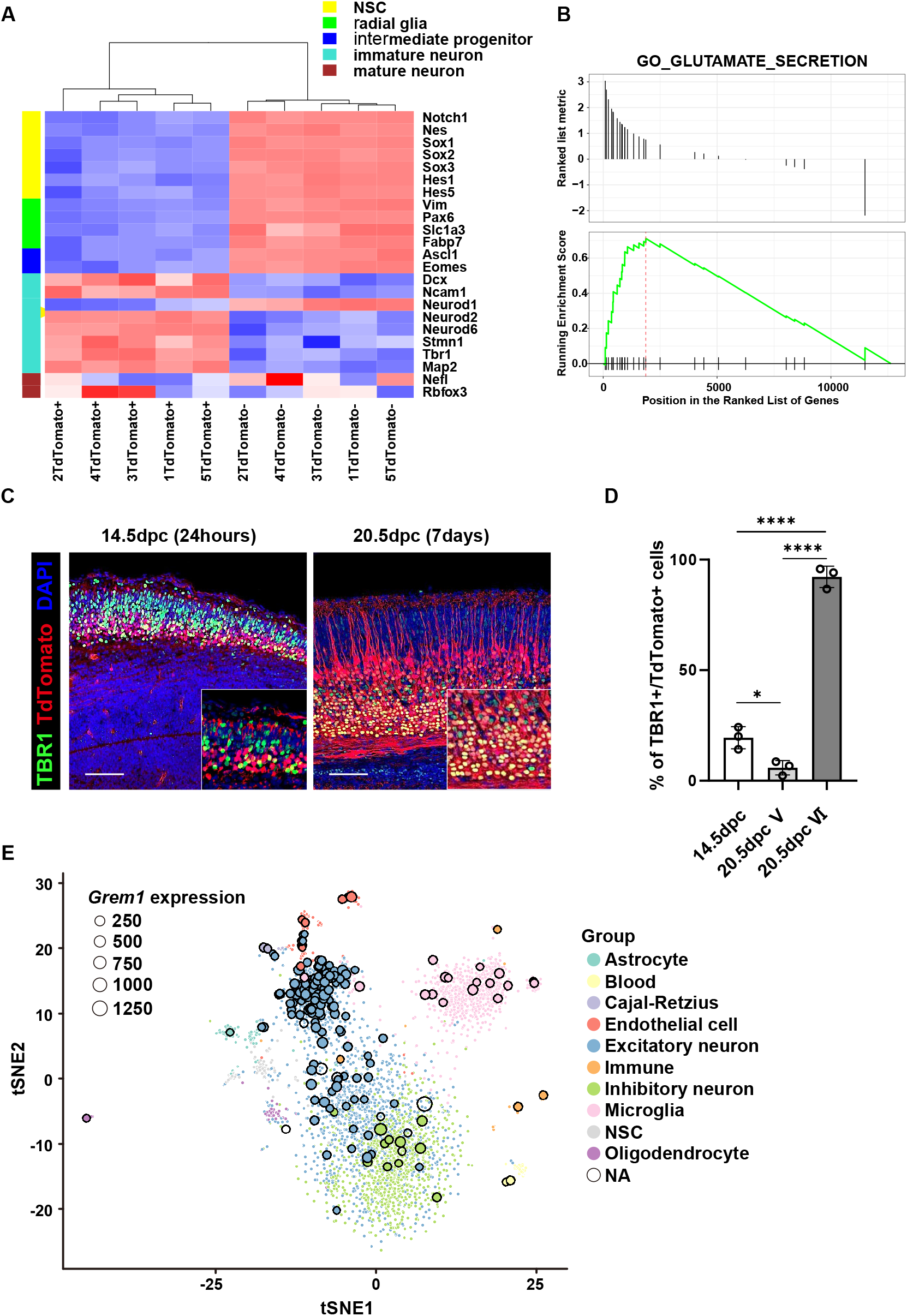
*Grem1* is expressed in the glutamatergic excitatory neuron lineage cells. (A) Heatmap depicting unsupervised clustering of TdTomato^+^ and TdTomato-cells isolated from 14.5dpc *Grem1reporter* (red) mice induced with tamoxifen at dpc13.5 based on expression of representative differentiation marker transcripts for neural stem cell (NSC), radial glia, intermediate progenitor, immature neuron, and mature neuron. (B) Gene set enrichment analysis (GSEA) for Glutamate Secretio<n genes between TdTomato+ and TdTomato-samples from (A). Normalised enrichment score (NES) = 2.16, p = 0.0049. (C) Representative images of immunofluorescence staining of 14.5 and 20.5 dpc telencephalon from *Grem1*reporter (red) mice induced with tamoxifen at dpc13.5, TBR1 (green), DAPI (blue). Scale bar = 100 μm. (D) Quantification of (C) showing the percentage of TdTomato^+^ cells that were also TBR1^+^ in 3 representative fields from 3 biological replicates. (E) tSNE plot of human scRNAseq dataset. *GREM1* expressing cells outlined in black. Dot size represents Grem1 expression value as indicated.

To assist with identification of the neuronal subtype likely generated from *Grem1-* expressing TdTomato^+^ cells, we also performed GSEA on DEG transcripts between TdTomato^+^ and TdTomato^-^ cells. We found a significant enrichment of a glutamate secretion gene set in the DEGs (NES = 2.16 p = 0.0049) (**Fig.3B**), whereas GABAergic and dopaminergic gene sets were not enriched (NES = −0.93 p = 0.75, NES = −0.69 p = 0.87 respectively) (**Fig.S4A**). Consistent with the GSEA, glutamatergic neuron markers, such as Slc17a7 (vGlut1), Grin1 and Grin2b were significantly upregulated in Grem1 expressing cells at 14.5 dpc using a candidate gene approach, whereas gabaergic and dopaminergic neuron markers, such as Slc6a1 (GABA transporter 1), Gad1, Gad2, and Th (Tyrosine hydroxylase) were significantly downregulated (p<0.05, **Fig.S4B**). Tbr1 plays a central role in the development of early-born cortical excitatory neurons and regulates the connectivity of layer VI neurons (Hevner et al., 2001). To confirm that Grem1 cells generate excitatory neuronal lineages, we undertook immunohistochemical staining with Tbr1. This revealed *Grem1*^-^ expressing TdTomato^+^ cells at 14.5 dpc expressed very low to no Tbr1, however they did give rise to Tbr1^+^ neurons in cortical layer VI at 20.5 dpc (**Fig.3C,D**).

To understand the relevance of our mouse focused study to the human setting, we reanalyzed publicly available human scRNAseq data generated from human brain at mid-gestation (22-23 weeks post-conception) (Fan et al., 2018). We categorized each human cell in the dataset as either *GREM1^+^* or *GREM1^-^* based on the presence or absence of *GREM1* RNAseq counts for each cell. Next, we compared the transcriptional profiles of our mouse embryonic brain *Grem1-*expressing TdTomato^+^ cells and TdTomato^-^ cells, with the human mid-gestational brain *GREM1*^+^ and *GREM1^-^* cells using a multidimensional scaling plot (**Fig.S4C**). Samples within the same group cluster tightly together, dimension 2 separates the samples along species lines, while dimension 1 clearly shows a similar separation of samples based on altered expression profiles depending on *Grem1/GREM1* status in mouse and human developing brain (**Fig.S4C**). By mapping the *GREM1*^+^ cells onto a tSNE plot generated from the human scRNAseq data, this revealed that most *GREM1*-expressing cells accumulated in the excitatory neuron cluster within the human mid-gestational cortex (**Fig.3E**). Lastly, we analysed the expression of BMP-signaling components in the human scRNAseq dataset (**Fig.S4D,S5**). Interestingly, of the pathway antagonists and similar to *GREM1, Sclerostin domain-containing protein 1 (SOSTDC1)* transcripts were also highly enriched in the excitatory neuron cluster, with *CHRD, NOG, BMP Binding Endothelial Regulator* (*BMPER*) and *Follistatin* (*FST)* transcripts also enriched in specific cell types (**Fig.S4D**). Transcripts encoding BMP2,4,7 that are antagonized by GREM1 were detected in various cell populations, however expression of *Bmp2,7* was inversely correlated with *Grem1* expression in the excitatory neuron cell population suggesting paracrine pathway regulation (**Fig.S5**). Expression of the BMP target genes, *Id1,3,4* was also inversely correlated with *Grem1*-expression in excitatory neurons, consistent with our developmental mouse brain data (Fig 2B). This highlights the complexity of regulation of BMP signaling during development of the cortex.

### Grem1 promotes proliferation and neural differentiation in neural stem / progenitor cells *ex vivo*

Next we collected NSPCs from embryonic brains of *Grem1^flox/flox^* mice at 14.5 dpc and transduced the cells ex vivo with control, *Cre*-expressing or *Grem1-* expressing lentivirus to generate control *Grem1^flox/flox^, Grem1^-/-^* and *Grem1* over-expressing (*Grem1^O/E^*) primary cultures. Endogenous Grem1 was detected in *Grem1^flox/flox^* control cultures by western blot, with absent or elevated protein levels in *Grem1^-/-^* and *Grem1^O/E^* cultures, respectively (**Fig.4A**). Control *Grem1^flox/flox^* cultures were responsive to BMP pathway induction as determined by increased BRE-luciferase reporter activity and expression of the BMP target genes, *Id1/2/3/4*, following addition of recombinant human BMP2 (**Fig.4B, Fig.S6A**). This induction of BRE-reporter activity and increase in BMP target gene transcript levels was significantly enhanced in *Grem1^-/-^* cells and attenuated in *Grem1^O/E^* cells (**Fig.4B, Fig.S6A**). This confirmed that Grem1 acts as an antagonist of BMP2 and suppresses downstream transcriptional targets in embryonic NSPCs.

**Fig. 4.**
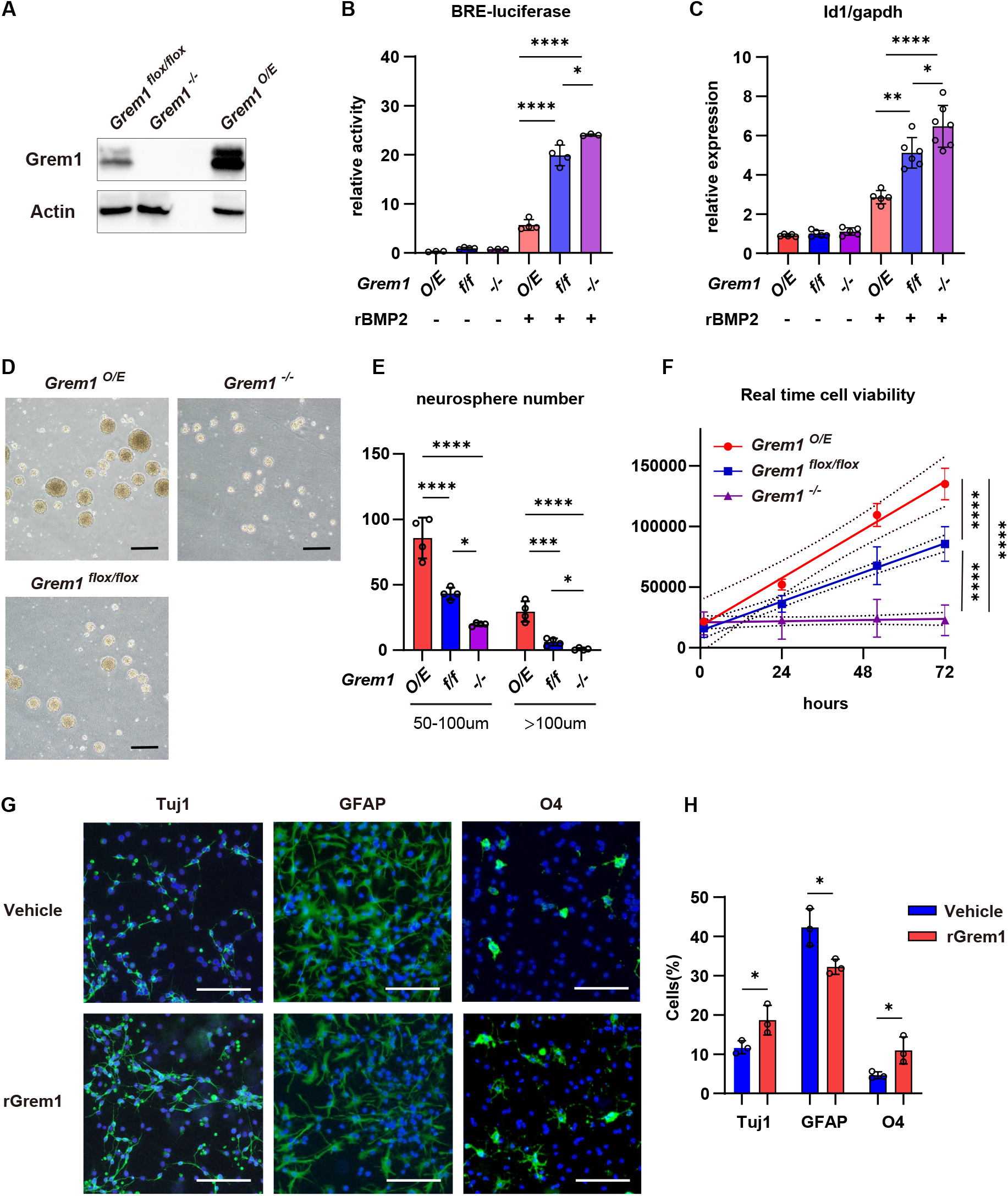
Gremlin1, a BMP antagonist, promotes proliferation and neuronal differentiation in neural stem / progenitor cells *ex vivo*. (A) Detection of Grem1 protein levels in control *Grem1^flox/flox^, Grem1^-/-^* and *Grem1^O/E^* NSPC by western blot. (B) BRE-reporter activity relative to internal control reporter. Data obtained from 4 independent experiments each performed in triplicate. (C) BMP target gene, *Id1*, mRNA expression normalized to *Gapdh* in *Grem1^flox/flox^, Grem1^-/-^* and *Grem1^O/E^* NSPC treated with vehicle or rBMP2 for 24h. Columns, mean; bars, SD. One way ANOVA with Tukey’s multiple test. Results from 5 independent experiments performed in triplicate. (D) Representative images of *Grem1^flox/flox^, Grem1^-/-^* and *Grem1^O/E^* neurosphere cultures. Scale bar = 100 μm. (E) Quantification of neurosphere number and size from (D) in 4 representative fields from 4 independent experiments. One way ANOVA with Tukey’s multiple test. (F) Cell viability normalized to 0h timepoint for each culture. Results from 3 independent experiments each performed in triplicate and analyzed using linear regression. (G) Representative images of immunofluorescence staining to identify Tuj 1^+^ neurons, GFAP^+^ astrocytes and O4^+^ oligodendrocytes in differentiated cultures from wild type NSPCs treated with vehicle or rGrem1. (H) Quantification of (G), showing the percentage of marker^+^ cells in 3 representative fields from 3 independent experiments. t-test. Scale bar = 100 μm. *p<0.05, **p<0.01, ***p<0.001, ****p<0.0001

We next employed neurosphere assays to determine the effect of Grem1 modulation on proliferation of embryonic NSPCs. Only mitogen responsive cells proliferate to form clusters termed neurospheres, where sphere size correlates with proliferative ability (Mori et al., 2006, Reynolds and Rietze, 2005). *Grem1^O/E^* cells form significantly more and larger neurospheres than control *Grem1^flox/flox^* cells, whereas *Grem1^-/-^* cells form significantly fewer (**Fig.4D,E**). To understand this phenomenon from the aspect of cell viability, we evaluated the proliferation rates of *Grem1^flox/flox^, Grem1^-/-^* and *Grem1^O/E^* cultures. Over-expression of *Grem1* significantly increased the rate of NSPC proliferation compared to control *Grem1^flox/flox^* NSPCs, while conversely the number of viable *Grem1^-/-^* cells did not increase over the 72h time course analysed (**Fig.4F**). These results suggested that Grem1 contributes to the proliferation of NSPCs. Following on from this observation, we wanted to assess the role of Grem1 in the differentiation of NSPCs to neurons, astrocytes and oligodendrocytes. When cultured in recombinant BMP2-containing differentiation media, addition of 1ng/ml recombinant GREM1 significantly increased the number of NSPCs that differentiated into Tuj1^+^ neurons and O4^+^ oligodendrocyte lineage cells, and decreased the number of GFAP^+^ astrocytes, in comparison to vehicle treated control NPSCs (**Fig.4G,H**). This suggests that Grem1 may regulate the differentiation potential of NPSCs.

### *Grem1* is required for normal cortical development

In order to assess the functional role of *Grem1* in mouse forebrain development in vivo, we generated tissue-specific *Grem1* conditional knockout mice using the *empty spiracles homeobox 1 (Emx1)-cre* driver. We first verified that *Emx1-cre* generates efficient cre-mediated recombination in the dorsal telencephalon at 14.5 dpc by visualizing TdTomato^+^ cells in reporter *Emx1-cre; Rosa26LSLTdtomato* mouse brains (**Fig.S7A**). Next we confirmed that *Emx1-cre; Grem1^flox/flox^* (*Grem1* conditional knockout, *Grem1^cKO^*) mice had a significant reduction in *Grem1* RNA in the developing forebrain compared with cre negative littermate controls by ISH at 14.5 dpc (**Fig.5A**) and real time RT-PCR at postnatal day0 (**Fig.5B**). *Grem1^cKO^* mice were viable and fertile. To examine the morphological consequences of conditional *Grem1* loss in the developing mouse brain we performed Nissl staining on tissue samples from *Grem1^cKO^* and cre negative littermate controls. At 10 weeks of age, total cortical thickness was significantly reduced in *Grem1^cKO^* mice in comparison with *Grem1^flox/flox^* littermate controls both in males and females, due to significantly thinner cortical layers V and VI (**Fig.5C, Fig.S7B**). The position and thickness of each layer was confirmed by immunohistochemistry using markers specific for each layer, i.e. CDP for layer II – IV, Ctip2 for layer V and VI and Foxp2 for layer VI (**Fig.5D,E**). Cellular density was also more sparse in *Grem1^cKO^* mice in comparison with *Grem1^flox/flox^* littermate controls (**Fig.S7C,D**). Conversely, the other predominant region of *Emx1-cre* driver activity, the hippocampus, that has important functions in memory for navigation pertinent to our behavioural testing, displayed normal morphology and cell density in *Grem1^cKO^* mice (**Fig.S7E**).

**Fig. 5.**
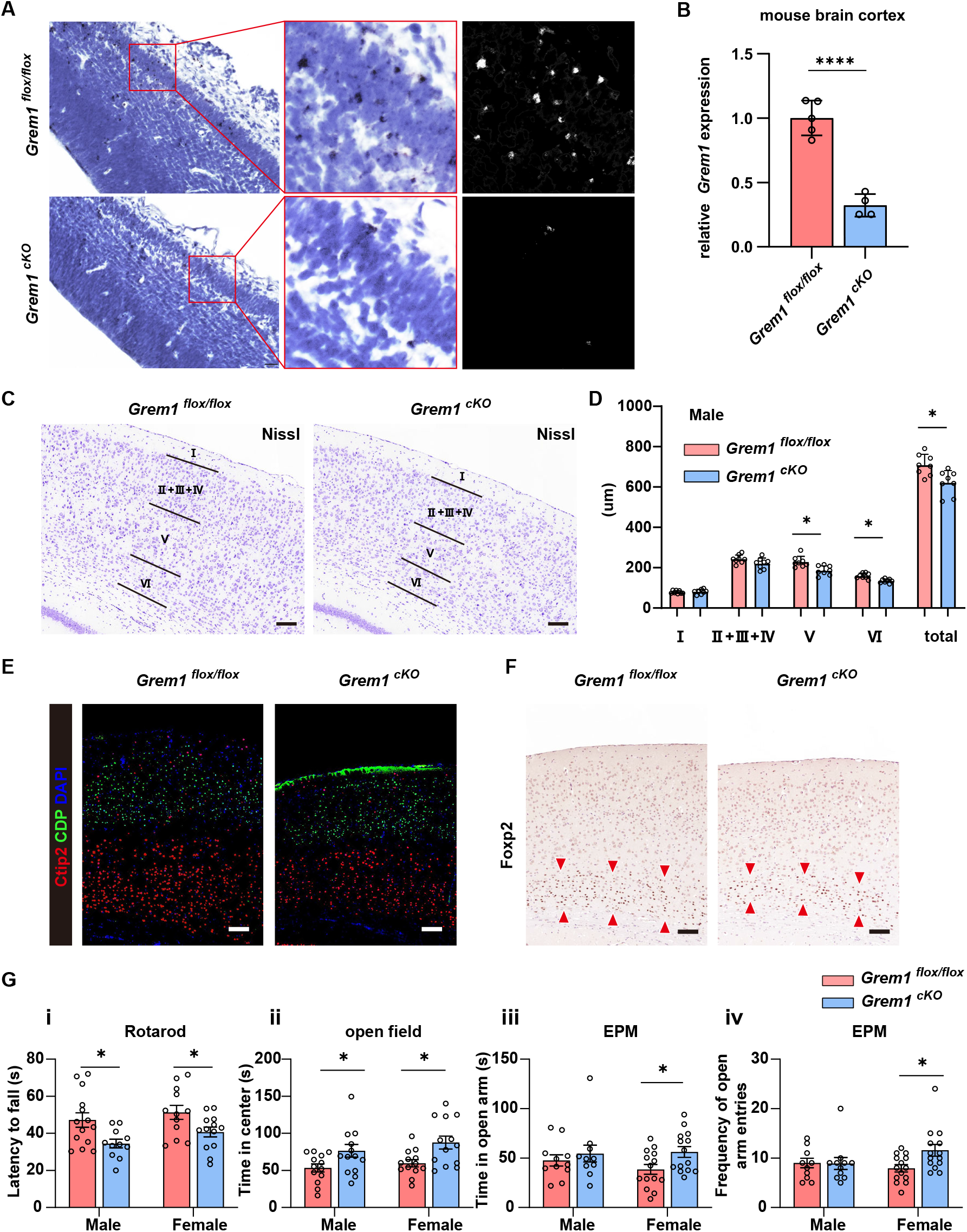
Cortical development, motor balance, and fear is impaired in Grem1 conditional knock out mice *Grem1* is required for normal cortical development. (A) Representative *Grem1* expression in the dorsal telencephalon of *Grem1^cKO^* mice and *Grem1^flox/flox^* littermate controls by ISH at 14.5 dpc. Monochrome images were prepared for quantitation of the *Grem1* ISH signal with ImageJ. n = 3 (B) *Grem1* mRNA levels normalized to *Gapdh* were determined using real time qRT-PCR in mouse cortical brain samples collected at postnatal day 0 from *Grem1^cKO^* mice and littermate controls n=4 biological replicates. Columns, mean; bars, SD. t-test. ****p<0.0001 (C) Representative histological images of cortical layers of *Grem1^cKO^* mice and *Grem1^flox/flox^* littermate controls using Nissl at 10 weeks of age. Scale bar = 100 μm. (D) Quantification of cortical layer thickness from (C) compared in 8 pairs of male *Grem1^cKO^* mice and littermate controls at 10 weeks of age. Female, n=7 control and n=7 *Grem1^cKO^*. t-test, *p<0.05. (E) Representative images of immunofluorescence staining of cortex of *Grem1^cKO^* mice and littermate controls at 10 weeks of age to visualise layer II – IV with marker CDP (green), layer V and VI with marker Ctip2 (red), and DAPI (blue). n = 3 Scale bar = 100 μm. (F) Representative images of immunohistochemical staining of cortex of *Grem1^cKO^* mice and littermate controls at 10 weeks of age to visualise layer VI marker Foxp2 IHC (shown in red arrowheads). n = 3 Scale bar = 100 μm. (G) Behavioral testing was performed to compare *Grem1^cKO^* and *Grem1^flox/flox^* mice using age and sex matched littermates at 7-10 weeks of age. Columns, mean; bars, SEM. (i) Rotarod test latency to fall. Male, n=14 control and n=11 *Grem1^cKO^*, Female, n=12 control and n=13 *Grem1^cKO^*. t-test, *p<0.05. (ii) Open field test, cumulative duration spent in the center area. Male, n=13 control and n=14 *Grem1^cKO^*, Female, n=13 control and n=13 *Grem1^cKO^*. t-test, *p<0.05. (iii, iv) Elevated plus maze test. (iii) Cumulative duration spent in open arms. (iv) The number of entries to open arms. Male, n=12 control and n=11 *Grem1^cKO^*, Female, n=14 control and n=14 *Grem1^cKO^*. t-test, *p<0.05.

Our earlier transcriptional network analysis from bulk RNAseq data identified an *Id1* associated gene cluster that was differentially regulated in the *Grem1-* expressing TdTomato^+^ cells from the cortex at 14.5 dpc (**Fig.2E**). From this cluster, *Lrrtm3* and *Ryr3* were selected for further analysis because of their known functions in synapse development (Balschun et al., 1999, Um et al., 2016, Roppongi et al., 2017, Futagi and Kitano, 2015, Linhoff et al., 2009). The expression of these two transcripts were significantly downregulated in the cortex of *Grem1^cKO^* mice compared with littermate controls at postnatal day 10 (**Fig.S7F**). This provides further evidence of the relationship between *Grem1* expression and these genes.

### Motor coordination and fear responses were impaired in *Grem1^cKO^* mice

To assess the functional consequences of embryonic *Grem1* deletion in the mouse forebrain, we undertook behavioural testing with *Grem1^cKO^* mice in comparison with *Grem1^flox/flox^* littermate controls. We compared motor balance using the Rotarod test. Latency to fall was significantly shorter in *Grem1^cKO^* mice both for males and females, suggesting an impaired motor balance with loss of *Grem1* (**Fig.5G**). During the open field test used to assess exploratory behaviours, both male and female *Grem1^cKO^* mice spent significantly more time in the central area away from the walls than littermate controls (**Fig.5G**). The total distance moved and velocity of movement were similar between *Grem1^cKO^* and littermate controls (data not shown). Behavioral testing using an elevated plus maze also indicated that female *Grem1^cKO^* mice spent significantly more time in the open maze arms and entered into the open arms more frequently than littermate controls (**Fig.5G**). Both of the exploratory behavior tests indicate that conditional loss of *Grem1* leads to reduced anxiety-like behavior. Lastly, we used the Y maze test, to assess the short-term memory of *Grem1^cKO^* and littermate controls. We observed no difference in the number of entries made to the novel arm of the Y maze between groups (**Fig.S8**), suggesting that *Grem1* expression is not required for short-term memory function.

## Discussion

The spatiotemporal regulation of BMP signaling in brain development is poorly understood. Here we focus on the expression and function of the BMP antagonist Grem1 during brain development, using transgenic lineage tracing, gene expression, in vitro culture and conditional knock out approaches.

In the developing mouse brain *Grem1* is first expressed at 13.5 dpc in the actively dividing dorsal telencephalon cells that retain mitotic potential (**Fig.1E,F**). *Grem1*-expression identifies neuroprogenitors committed to the NeuN+ layer V and Tbr1+ VI neurons. Transcriptomic profiling using our mRNAseq data from embryonic *Grem1*-reporter mouse brains and publicly available human developmental brain scRNAseq data suggests that *Grem1/GREM1* expression is primarily associated with markers of glutamatergic excitatory neuronal lineages, rather than GABAergic or dopaminergic neurons or other cell types of the developing brain (**Fig.3B,E**, **Fig.S4A,B**). Consistent with this, *Grem1* was not expressed in the ganglionic eminences, from which GABAergic interneurons originate; nor were GABAergic transcripts upregulated or enriched by GSEA in *Grem1*-expressing TdTomato^+^ cells in the dorsal telencephalon using bulk RNAseq analysis (**Fig.S4A, B**). Likewise Grem1-lineage traced cortical neurons expressed the excitatory marker, Tbr1 (**Fig.3C**). *Grem1*-expression labels and generates important excitatory neuronal lineages, i.e. pyramidal neurons, that play high-level cognitive functions in the neocortex. As Grem1 expression is required for the correct development of early-born excitatory neurons of the neocortex (**Fig.1G**), misregulation of BMP signaling likely affects the balance between excitatory and inhibitory neuronal activities, which has been implicated in neurodevelopmental disorders (Dani et al., 2005).

Our analysis of scRNAseq data from human midgestational cortex (22-23 weeks post-conception) suggests that there is coordinated regulation of BMP agonists and antagonists at this developmental stage in humans, but that there may be also be functional redundancies (**Fig.S4D**). Noggin is the most extensively studied of the BMP antagonists in the CNS, where it is a recognized neural inducer during gastrulation (Lamb et al., 1993). Addition of recombinant Noggin to neural stem cells promotes neuronal and suppresses astrocytic differentiation *in vitro* (Mikawa and Sato, 2011). In our study, we confirmed that Grem1 acts as a BMP antagonist in embryonic NSPC cultures, suppressing downstream BMP transcriptional targets (**Fig.4,S6**). Similar to Noggin, we determined that Grem1 promotes proliferation and neural differentiation, at the expense of astrocytic lineages *in vitro* (**Fig.4**). BMP transcriptional targets were significantly downregulated in *Grem1-*expressing TdTomato^+^ cells also in vivo (**Fig.2B**) and *Grem1^+^* excitatory neurons in the human embryonic cortex (**Fig.S5**). Together these results suggest that Grem1 expressing cells are fated to become neurons via antagonism of potential paracrine BMP signals (**Fig.S4D,S5**) (Furuta et al., 1997, Imura et al., 2008, Choe and Pleasure, 2018).

Our histological analyses identified that the cortex (particularly neocortical layers V and VI) of *Grem1^cKO^* animals is significantly thinner, with less cells, in comparison to littermate controls, with no obvious morphological changes in the hippocampus. Our in vitro experiments confirmed that Grem1 drives proliferation and neuronal differentiation in NSPC, thus this cortical phenotype is likely due to reduced proliferation of embryonic NSPCs in *Grem1^cKO^* mice. Layer V neurons target the spinal cord, cerebellum, striatum and the thalamus (Shipp, 2007) and play roles in movement preparation, movement guidance, and the execution of well-timed movements (Baker et al., 2018, Li et al., 2015). The observed decrease in layer V neurons would lead to impaired preparation for and coordination of movement in *Grem1^cKO^* animals and explain the motor balance defect in these animals in comparison to litter mate controls. Emerging evidence suggests that layer VI neurons play a central role in modulating thalamic and cortical neurons during sensory processing (Wang et al., 2016). Our behavioural testing confirmed that loss of Grem1 expression significantly impaired fear sensitivity in *Grem1^cKO^* animals, compared to littermate controls, consistent with a deficiency in layer VI neurons with roles in sensory connections. BMP signaling is known to promote synaptogenesis (Xiao et al., 2013, Shen et al., 2004) and altered synaptogenesis can affect both fear sensitivity and motor abilities (Ha et al., 2016, Wood and Shepherd, 2010). Network analysis of DEG between *Grem1-*expressing TdTomato^+^ and TdTomato^-^ cells from the developing mouse dorsal telencephalon identified a gene cluster of inter-related transcripts in which the BMP target gene, *Id1*, acts as a hub. Genes in the cluster play roles in neuron morphogenesis, synapse and axon maturation (**Fig.2D,E**). For example, *Leucine-rich-repeat transmembrane neuronal proteins 3/4* (*Lrrtm3/4*) genes encode synapse organizing proteins critical for the development and function of excitatory synapses (Um et al., 2016, Roppongi et al., 2017, Linhoff et al., 2009). Likewise, the *Ryanodine receptor3(Ryr3* gene encodes a member of a family of receptors that shape synaptic transmissions(Balschun et al., 1999) by amplifying spike-driven calcium signals in presynaptic terminals, and consequently enhancing the efficacy of transmitter release (Futagi and Kitano, 2015). In *Grem1^cKO^* animals, the mRNA expression of *Lrrtm3* and *Ryr3* was significantly decreased in comparison to littermate controls (**Fig.S7F**). These results suggest *Grem1* may contribute to synapse formation and function by inhibiting BMP signaling and expression of associated gene networks, the subject of further studies.

In summary, this is the first study to reveal the important function of Grem1 in cortical development, particularly as a marker, and maker, of excitatory neuroprogenitors. In the future, Grem1 may hold value beyond understanding the cellular biology of brain development and function as we develop new approaches to help tackle complex neurodevelopmental and neurological diseases.

## Materials and Methods

### Mice

*Grem1-CreERT* transgenic mice (Worthley et al., 2015) were crossed with *R26-LSLTdTomato* mice (B6.Cg-*Gt(ROSA)26Sor^tm14(CAG-tdTomato)Hze^*/J, JAX 007914) to generate a tamoxifen-induced *Grem1* reporter line. Pregnant dams were administered 6mg of tamoxifen by oral gavage to induce embryonic *Grem1*-tracing, with some dams bearing 13.5 dpc embryos also intraperitoneally injected with 5-Bromodeoxyuridine (BrdU, Roche, 40 mg/kg) to track cell division. *Grem1^flox/flox^* mice (Gazzerro et al., 2007) were crossed with *Emx1-cre* mice (B6.129S2-*Emx1^tm1(cre)Krj^*/J, JAX 005628) to generate *Emx1-cre* mediated *Grem1* conditional knockout mice *(Emx1-cK(O)* The line was maintained by crossing *Emx1^-cre/+^; Grem1^flox/flox^* mice with *Grem1^flox/flox^* mice.

All mice were on the C57BL6/J background and experimentation was conducted following approval by SAHMRI Animal Ethics Committee (approval number SAM284) in accordance with the Australian code for the care and use of animals for scientific purposes, 8^th^ edition.

### Preparation of single cell suspensions and flow cytometry

Pregnant *Grem1creERT; Rosa26LSLTdtomato* mice were administered tamoxifen at 13.5 dpc. At 14.5 dpc, dorsal telencephalons of the embryos were dissected in cold phosphate-buffered saline (PBS) and the meningeal membranes were removed. Neurocult Enzymatic Dissociation kit (Stemcell Technologies) was used for cell dissociation according to the manufacturer’s protocol. Dissociated cells were resuspended in FACS buffer containing DAPI (0.5μg/ml). Sorting and analyses were carried out on a FACS Fusion flow cytometer (Becton-Dickinson). Dead cells were excluded by gating on forward and side scatter and by eliminating DAPI-positive events. The cells harvested from *cre* negative littermate control mice were used to set background fluorescence levels. Viable cells were sorted into FACS buffer, collected via centrifugation and resuspended in Trizol (Invitrogen) for RNA extraction.

### RNA extraction and mRNA seq

RNA was extracted from sorted cells in Trizol according to the manufacturer’s protocol with the exception of addition of glycogen (20μg/μl) and isopropanol precipitation overnight at −80° C to maximize yield. RNA quality and quantity were analyzed using a NanoDrop and TapeStation. Total RNA was converted to strandspecific Illumina compatible sequencing libraries using the Nugen Universal Plus mRNA-Seq library kit from Tecan (Mannedorf) as per the manufacturer’s instructions (MO1442 v2). Briefly, 500ng of total RNA was polyA selected and the mRNA fragmented prior to reverse transcription and second strand cDNA synthesis using dUTP. The resultant cDNA was end repaired before the ligation of Illumina– compatible barcoded sequencing adapters. The cDNA libraries were strand selected and PCR amplified for 12 cycles prior to assessment by Agilent Tapestation for quality and Qubit fluorescence assay for quantity. Sequencing pools were generated by mixing equimolar amounts of compatible sample libraries based on the Qubit measurements. Sequencing of the library pool was performed with an Illumina Nextseq 500 using single read 75bp (v2.0) sequencing chemistry.

### Bioinformatic analysis

#### RNAseq data processing

Fastq files from the sequencing run were firstly subjected to quality controls with FastQC version 0.11.3. Raw reads with low quality were removed using Trim Galore alignment (Phred score less than 28 and/or reads contains adaptor sequences). After trimming, all bases with low quality scores and adaptor sequences were removed. The trimmed reads were mapped to Ensembl mouse genome (GRCm38) with STAR 2.4.2a. No more than 1 base mismatch was allowed. Only uniquely mapped reads were retained. Option -quantMode was enabled to generate gene level quantification. The counts files were then merged into an expression table for downstream differential expression analysis.

#### Differential expression analysis

Differentially expressed genes (DEG) between the *Tdtomato*^+^ (*Grem1* positive) and *Tdtomato*-(*Grem1* negative) populations were analyzed with edgeR packages in R version 3.6.0 (Robinson et al., 2010). Reads were inspected through multidimensional scaling plot and outliers were removed. TMM normalized counts were log2-transformed and counts per million (CPM) obtained. Paired comparisons between the *Tdtomato*^+^ (*Grem1* positive) and *Tdtomato*-(*Grem1* negative) populations were performed. DEG reported as significant were selected by requiring both adjusted P value (FDR) ≤ 0.05 and absolute value of log2 fold change ≥ 2.0.

#### Supervised Weighted Gene Correlation Network Analysis (WGCNA)

Only DEGs were used for subsequent network analysis. The gene network was constructed using the R package WGCNA following the procedure as described (Langfelder and Horvath, 2008). After the low expression genes (FPKM < 1) had been filtered out from all gene expression libraries, Pearson’s correlation based adjacency was calculated on the basis of pairwise correlations of gene expression within *Tdtomato*^+^ (*Grem1* positive) samples. Topological overlap of the correlations was used to weight the edges of the correlation network. The higher the weight, the stronger the interaction between two genes. Connectivity for a single gene was calculated as the sum of weights relative to the rest of the genes, and the top 5% of genes with the highest connectivity in the network defined as hub genes. For visualisation, a heat map was generated using the TOMplot() function in WGCNA R package (version 1.68), with dissimilarity topological overlap (1-topological overlap), employed for hierarchical clustering. To generate the network plot, the weights of the network were cut off at 0.1. Hypergeometric enrichment tests for each module-defined gene list were performed with the enricher() function, as part of clusterProfiler R package version 3.13.0 (Yu et al., 2012). Bonferroni adjustment for p value was used, only gene sets with an adjusted p value less than 0.05 were considered for interpretation of the biological function modules.

#### Gene set enrichment analysis (GSEA)

To utilize the Molecular signatures database (MSigDB), we accessed all gene sets (.gmt file, version 6.2) from the Broad Institute and chose a subset for further analyses including BioCarta, Hallmark, Gene Ontology (GO), Kyoto Encyclopedia of Genes and Genomes (KEGG), Pathway Interaction Database, and the Reactome Pathway Database. To enable cross species comparisons, mouse gene ensembl IDs were converted to human orthologous gene symbols using biomaRt R package, version 2.41.7 (Durinck et al., 2009). Humanised gene lists for the whole transcriptome were ranked based on logfc values by comparing TdTomato^+^ and TdTomato-cells. The R package clusterProfiler version 3.13.0 (Yu et al., 2012) was used to scan through all the gene sets mentioned above, using Benjamini & Hochberg adjusted p values.

#### Analysis of publicly available single cell RNA seq dataset

Single cell RNAseq raw expression data was accessed for human mid-gestational brain (22 and 23 weeks post-conception) cortex samples from GSE103723 (Fan et al., 2018). The cells from two individual samples were grouped together, with the sum of reads calculated for each gene to represent the data structure of bulky transcriptomics. The limma package(Ritchie et al., 2015) was used to remove batch effects with removeBatchEffect() and to generate multidimensional scaling plots. To understand the expression pattern of human *GREM1* and its possible contribution to neural differentiation, normalized counts and tSNE coordinates were employed for visualization using plots generated with ggplot2 package in R environment (Wickham, 2009). To investigate cell types expressing BMP antagonists, pearson’s Chi-Square test of independence was performed in the R environment with chisq.test() function. Testing variables were gene expression categorised into high and low by median expression, and using cell type identifiers from the original scRNAseq study (Fan et al., 2018). Scaled standardised residuals for each gene were used to plot the heatmap with ComplexHeatmap package (Gu et al., 2016).

#### *In situ* hybridization (ISH)

All ISH analyses were performed on formalin-fixed and paraffin-embedded mouse tissue samples using RNAscope technology (RNAscope 2.5 HD Detection Kit, RNAscope Multiplex Fluorescent Reagent Kit v2, Advanced Cell Diagnostics) following the manufacturer’s instructions. Briefly, tissue sections were baked in a dry oven (HybEZ II Hybridization System, Advanced Cell Diagnostics) at 60°C for 1 h and deparaffinized, followed by incubation with an H_2_O_2_ solution (Pretreat 1 buffer) for 10 min at RT. Slides were boiled in a target retrieval solution (Pretreat 2 buffer) for 15 min, followed by incubation with a protease solution (Pretreat 3 buffer) for 30 min at 40°C. Slides were incubated with a mouse *Grem1* probe (NM_011824.4, region 398-1359, catalogue number 314741) for 2 h at 40°C, followed by successive incubations with signal amplification reagents. Staining was visualized with DAB or TSA Plus Cyanine 5 (NEL745001KT, PerkinElmer; 1:750). Combined ISH and immunofluorescence (IF)/immunohistochemistry (IHC) was undertaken by first performing ISH, followed by IF/IHC. For IF following ISH, the sections were blocked with blocking buffer (X0909, Dako) and then incubated with a primary antibody (rabbit polyclonal anti-RFP, Rockland, 600-401-379) overnight at 4°C. The sections were washed in 1x T-PBS 3 times and then incubated with Alexa Fluor 555-conjugated secondary antibody (Thermo Fisher Scientific) for 60 min at room temperature. The sections were mounted with ProLong Gold antifade reagent containing 4’6-diamidino-2-phenylindole (DAPI; Thermo Fisher Scientific), and fluorescence was examined using an inverse immunofluorescence microscope BZ-X710 (Keyence, Japan) with optical sectioning. For IHC following ISH, after inactivation of endogenous alkaline phosphatase with alkaline phosphatase blocking solution for 10 minutes (SP-6000, Vector Laboratories), the slides were blocked with blocking buffer (X0909, Dako) and incubated with a primary antibody (rabbit polyclonal anti-RFP, Rockland, 600-401-379) at 4 °C overnight. The sections were washed with PBS three times, followed by incubation with a horse anti-rabbit IgG polymer (MP-5401, Vector Laboratories) for 30 min at room temperature. The slides were washed with TBS three times, and signal developed using an alkaline phosphatase substrate (SK-5105, Vector Laboratories). The reaction was stopped by washing in TBS four times, and slides were counterstained with hematoxylin, dehydrated in EtOH, cleared in xylene, and coverslipped. Cells expressing more than 1 ISH signal were regarded as positive for Grem1 RNA.

#### Real time RT-PCR

RNeasy mini kits (Qiagen) were used to isolate RNA from snap frozen mouse brain tissues. RNA was reverse-transcribed into cDNA with cDNA master (Sigma). PCR amplification was performed with Kappa Sybr qPCR mix or Kappa Probe qPCR mix using QuantStudio7 (Applied Biosystems). The mouse GAPDH gene was used as an endogenous control. The following primers were used:

TaqMan probes and primers, Gremlin1 (IDT, Mm.PT.58.11631114), Id1 (IDT,Mm.PT.58.6622645.g), Id2 (IDT, Mm.PT.58.13116812.g), Id3 (IDT, Mm.PT.58.29482466.g), Id4 (IDT, Mm.PT.58.6851535)

Sybr primers, Gapdh-forward AAGGTCATCCCAGAGCTGAA, Gapdh-reverse CTGCTTCACCACCTTCTTGA Ryr3-forward TGCTGTCGCTTCCTTTGCTA, Ryr3-reverse CATCGATGGGGACGCTAGAC, Lrrtm3-forward TAGCAAATCAGGCTCCAGGG, Lrrtm3-reverse GAGTTCATGATGGACCCCACA

#### Immunohistochemistry

To collect 14.5 dpc, 17.5 dpc and 20.5 dpc embryonic brains, euthanized pregnant dams immediately underwent cardiac perfusion with 4% PFA. Whole embryos at 14.5 dpc, or dissected embryonic brains for 17.5 and 20.5 dpc were fixed in 4% PFA at 4°C overnight. Embryos and brains were then cryoprotected in 30% sucrose and frozen in OCT embedding medium. 16 μm sections were cut using a Leica CM1900 cryostat. To make paraffin sections, samples were post fixed in 10% neutral buffered formalin overnight and processed. 5 μm sections were cut using a Leica HM325 microtome. Sections were blocked with Protein Block Serum-Free (Dako) for 1 hour at room temperature, incubated overnight with first antibody at 4 °C, secondary antibody for 1 hour at room temperature, and coverslipped with Vectashield Antifade Mounting Medium (Vector laboratories). The following antibodies were used: Anti-TBRl(Abcam, ab183032, 1:400), Anti-NeuN (Abcam, ab104225, 1:500), Anti-β III tubulin (Sigma, T5076, 1:400), Anti-O4 (R & D systems, MAB1326, 1:500), Anti-GFAP (Dako, Z0334, 1:250), Anti-FOXP2 (Abcam, ab16046, 1:10000), Anti-Ctip2 (Abcam, ab18465, 1:800), Anti-CDP (Santa Cruz, sc-13024, 1:400), Anti-BrdU (Abcam, ab6326, 1:600), Anti-RFP (Rockland, 600-401-379, 1:1000), Anti-Sox2 (Millipore, AB5603, 1:400), Anti-Pax6 (Millipore, AB2237, 1:100) Images were acquired on a Leica SP5 spectral scanning confocal microscope.

#### Neural stem cell and progenitor cell culture

Before seeding cells, tissue culture dishes were coated with PDL (100μg/ml) and laminin (10μg/ml). Embryonic neural stem and progenitor cells were isolated from pregnant mice at 14.5 dpc. Dorsal telencephalons were dissected from each embryo in PBS, meningeal membranes removed and the tissue triturated to a single cell suspension. Cells were cultured in NeuroCult™ Proliferation Medium containing 20 ng/ml epidermal growth factor (EGF) (Stemcell Technologies). For differentiation assays, cells were seeded onto 4 well chamber slides (Thermofisher #NUN 177399) in Neurocult Differentiation medium (Stemcell Technologies). To induce recombination or overexpression of *Grem1*, cells collected from Grem1^flox/flox^ embryos were infected with plenti-EF1-*Cre-2a-sfGFP-2a-puro*, plenti-EF1-*Grem1-2a-sfGFP-2a-puro*, or plenti-EF1-*2a-sfGFP-2a-puro* lentivirus, and transduced cells were selected using puromycin for 5 days and used for experiments before passage 5. Lentivirus plasmids, psPAX2 and MD2.G were transfected to 293T cells to generate lentivirus, and viral supernatant was concentrated using Amicon-Ultra 100k spin columns. To assess neurosphere forming ability, cells were seeded in 6-well uncoated plates (1.6×10^5^ cells per well) and cultured for 5 days. The number of neurospheres sized 50-100 μm and >100 μm was counted. For cell viability assays, neural stem / progenitor cells were seeded in 96-well coated plates (1×10^4^ cells per well) and RealTime-Glo^TM^ MT Cell Viability Assay (Promega) was used with the continuous-read protocol at 0, 24, 52 and 72 hours. Recombinant human BMP2 (Prospec) was added to the differentiation medium.

#### Luciferase assay

BMP pathway activity was measured using the BMP response element luciferase reporter pGL3 BRE luciferase (addgene #45126), internal control pRL/TK-luciferase reporter and the dual luciferase reporter assay kit (Promega). Neural stem / progenitor cells were seeded in coated 24-well plates (1.6×10^5^ cells per well) and transfected with the reporter plasmids using xTremeGENE HP DNA transfection reagent (Roche). Cells were collected 48hr after transfection using Passive lysis buffer (Promega). The pGL3 empty vector was used as a control for BMP-independent changes in reporter activity.

#### Western blot

Cell lysates were solubilized with M-PER Mammalian Protein Extraction Reagent containing complete protease and phosphatase inhibitors. Lysates were separated by SDSPAGE and transferred to PVDF membranes (Millipore). After blocking with 5% nonfat skim milk in PBST for 30 min at room temperature, the membranes were incubated overnight at 4°C with anti-Gremlin1 antibody (R&D systems, AF956, 1:1000) or anti-βActin (Santa Cruz, sc-47778, 1:1000) in 0.5% nonfat skim milk in PBST. Membranes were washed with PBST and incubated with alkaline phosphatase-conjugated secondary antibodies (anti-rabbit IgG and anti-mouse IgG, GE healthcare life sciences, 1:10000) for 1 hr at room temperature. Finally, the blot was visualized with Immobilon HRP substrate (Millipore) using a Chemi Doc XRS1 (Bio-Rad).

#### Behavioral tests

Mice were submitted to Rotarod, open field, elevated plus maze, and Y maze tests at the age of 7-10 weeks. Mice while in their home cages were acclimatized to the behavior suite for at least 30 min prior to testing. The data was acquired blindly to the genotype.

Rotarod. Animals were placed on the rotarod (Panlab/Harvard Apparatus) that linearly increased rotation speed from 4 to 40 rpm during a 120 second period. An accelerating protocol was employed to eliminate the need for habituation to the rotarod. This procedure was repeated for a total of three trials per mouse, separated by 15min inter-trial intervals and the mean latency to fall from the rotarod in seconds was compared between wild type littermate controls and *Grem1^cKO^* mice to assess motor coordination.

Open field test. The open field test was conducted in four identical square arenas (50×50×50cm) surrounded by walls. Mice were individually placed in a corner of a clean arena and allowed to explore for 10 min. For the purpose of data collection, the arena was conceptually partitioned into two zones: a virtual center zone of 23×23 cm and a peripheral zone occupying the remaining area. Lower percentages of time spent in the center zone was used to indicate a higher level of anxiety.

Elevated plus maze test. The elevated plus maze consisted of a central square (8 × 8 cm) and four arms (29 cm long × 8 cm wide, two open arms with no railing and two closed arms enclosed by a transverse wall 20 cm in height). A mouse was placed in the center of the central square facing the open arm and allowed to explore the maze apparatus for 10 min. The time spent in any of the open arms was recorded and used as a measure of anxiety.

Y maze test. The apparatus consisted of three arms with an angle of 120° between each of the two arms. Each arm was 40 cm long × 8 cm wide × 15 cm high. Visual cues were placed on the walls of the mazes. The Y-maze test consisted of two trials separated by an inter-trial interval (ITI) to assess spatial recognition memory. The first trial (training) had a 5 min duration and allowed the mouse to explore only two arms of the maze, with the third arm (novel arm) being blocked. After 30min ITI, the second trial was conducted, during which all three arms were accessible for 5 min. Trials were recorded using a ceiling-mounted camera and analyzed by a video analyzer (Ethovision XT, Noldus) to determine the period that each mouse spent in each arm of the maze.

#### Statistics

All statistical analyses were performed using Graphpad prism 8, with methods and values summarised in each figure legend.

## Acknowledgements

We thank Dr. Randall Grose (SAHMRI, Australia) for flow cytometry assistance; Dr Mark van der Hoek and the Genomics Core (SAHMRI, Australia) for RNA-seq library preparation and sequencing, Dr. Hiroaki Fujiwara (The Institute for Adult Diseases, Asahi Life Foundation, Tokyo, Japan) for reviewing the manuscript; Regeneron and Prof Andrew Zannettino for providing *Grem1^flox/flox^*; *Emx1-cre* mice.

## Data availability

The accession number for all sequencing data reported in this paper is GEO: GSEXXXXXX.

---

**Fig.S1 *Grem1* starts to express after 12.5 dpc and is dramatically decreased after birth**

(A) Representative coronal images of immunofluorescence staining of 12.5 dpc telencephalon from *Grem 1*-reporter (red) mice induced with tamoxifen at 11.5 dpc. Scale bar = 100 μm. LV, lateral ventricle; NCx, neocortex

(B) *Grem1* mRNA levels normalized to *Gapdh* were determined using real time qRT-PCR in mouse cortical brain samples collected at p0, p10 and 4 weeks post-birth (n=4-6 mice/group, PCR conducted in triplicate) Columns, mean; bars, SD. One way ANOVA with Tukey’s multiple test. ****p<0.0001

**Fig.S2 Clustering of differentially expressed genes**

A heat map was generated to visualise the unsupervised hierarchical clustering of correlation scores for gene expression of DEG in TdTomato^+^ cells. Yellow colour indicates closely related and red not related. The color bar between the hierarchical clustering tree and plotting fields denotes the module membership of each gene.

**Fig.S3 *Grem1*-expressing cells are radial glial stem cell markers, Sox2 and Pax6 negative**

Representative images of immunofluorescence staining of 14.5 dpc neocortex from *Grem1-reporter* (red) mice induced with tamoxifen at dpc13.5, (A) Sox2 (green), DAPI (blue). (B) Pax6 (green), DAPI (blue). Scale bar = 100 μm.

**Fig.S4 Extended transcriptome analysis of bulk mRNAseq data from *Grem1-expressing* TdTomato^+^ and TdTomato^-^ telencephalon cells at 14.5dpc**

(A) Gene set enrichment analysis (GSEA) for genes involved in the regulation of (i) GABAergic synaptic transmissions and (ii) dopamine secretion that were differentially regulated between *Grem1-expressing* TdTomato^+^ and TdTomato^-^ cells. Normalised enrichment score (NES) = −0.93, p = 0.75 and NES = −0.69, p = 0.87.

(B) Unsupervised clustering of *Grem1-expressing* TdTomato^+^ and TdTomato samples based on expression of representative markers for glutamatergic, GABAergic, and Dopaminergic neurons, in bulk mRNAseq data. Marker transcript expression is displayed as a heat map with high expression in red to low expression in blue.

(C) Multidimensional scaling plot comparison of transcriptome of TdTomato^+^ and TdTomato^-^ cells isolated from 14.5dpc *Gremlreporter* mice induced with tamoxifen at dpc13.5 and *GREM1*^+^ and *GREM1*^-^ cells from scRNAseq of human mid-gestational cortex.

(D) GREM1 expression is associated with the excitatory neuronal lineage in the developing human cortex. Unsupervised clustering of human developmental brain cell populations from scRNAseq GSE103723 based on expression of lineage markers and transcripts encoding BMP antagonists *(FST, NOG, CHRD, BMPER, GREM1, SOSTDC1),* and BMP target genes *(ID1-4)*. Cell type is indicated by colour legend as per tSNE plot shown in Figure 4E. Chi-square testing was used to evaluate the level of each transcript in each cell population and the resulting standardized residual values are depicted in the heat map.

**Fig.S5 Grem1 suppresses intrinsic BMP signaling in developing human excitatory neurons, likely via antagonism of paracrine BMP ligands.**

(A) Heat map depicting expression of BMP ligands in human developmental brain cell populations from scRNAseq GSE103723. Cell types as per tSNE plot shown in Figure 4E. Chi-square testing was used to evaluate the level of each transcript in each cell population and the resulting standardized residual values are depicted in the heat map.

(B) GREM1 expression inversely correlates with enrichment of BMP2,7 and BMP target genes ID1, 3, 4 expression in human excitatory neurons in mid-gestation. Expression of transcripts encoding BMP ligands antagonised by GREM1 and ID target genes in *GREM1*^+^ and *GREM1^-^* cells is displayed from human scRNAseq GSE103723. One way ANOVA with Tukey’s multiple test. ns p>0.05, *p<0.05, **p<0.01,****p<0.0001

**Fig.S6 Grem1 acts to antagonize BMP signaling in NSPCs resulting in altered downstream transcript levels**

Transcript levels of BMP target genes, *Id2*, *Id3* and *Id4,* normalized to *Gapdh* in *Grem1^flox/flox^, Grem1^-/-^* and *Grem1^O/E^* NSPC treated with vehicle or rBMP2 for 24h.

Results from 5 independent experiments performed in triplicate. Columns, mean; bars, SD. One way ANOVA with Tukey’s multiple test. *p<0.05, ***p<0.001

**Fig.S7 Extended morphological, immunostaining and transcript analysis of *Grem1^cKO^* mice and littermate controls**

(A) Representative coronal section of TdTomato^+^ (red) cells in the dorsal telencephalon at 14.5dpc in *Emx1-cre; Rosa26LSLTdtomato* mouse brain. This cre-driver is broadly active across the telencephalon region in which *Grem1* is highly expressed. Scale bar = 100 μm. NCx, neocortex; H, hippocampus; Str, striatum; T, thalamus

(B) Quantification of cortical layer thickness compared between age and sex matched littermates at 10 weeks of age. Female, n=7 control and n=7 *Grem1^cKO^,.* t-test, *p<0.05.

(C) Representative histological images of neocortical layer 5 and 6 from *Grem1^cKO^* mice and *Grem1^flox/flox^* littermate controls at 10 weeks of age using Nissl. Scale bar = 20um.

(D) Quantification of (C) showing the cell number per area of layer V and VI using Nissl in 2 HPF of 3 biological replicates. Columns, mean; bars, SD. t-test. *p<0.05. Scale bar = 20um.

(E) Representative histological images of hippocampus from *Grem1^cKO^* mice and *Grem1^flox/flox^* littermate controls at 10 weeks of age using Nissl. 8 pairs of males and 7 pairs of females were analyzed. Scale bar = 200 μm.

(F) Transcripts from the *Id1* associated gene cluster (Fig 3E), *Ryr3* and *Lrrtm3* were evaluated in *Grem1^cKO^* and *Grem1^flox/flox^* littermate control cortex at p10 using real time qRT-PCR. *Ryr3* and *Lrrtm3*levels were normalized to *Gapdh.* n=4-6 biological replicates. Columns, mean; bars, SD. t-test. *p<0.05, ****p<0.0001

**Fig.S8 Memory is not impaired in *Grem1^cKO^* mice compared to *Grem1^flox/flox^* littermate controls**

(A,B) Y maze test. Behavior was compared between age and sex matched *Grem1^cKO^* mice and *Grem1^flox/flox^* littermate controls at 7-10 weeks of age. Male, n=12 control and n=10 *Grem1^cKO^*, Female, n=14 control and n=13 *Grem1^cKO^*. t-test. (A) Cumulative duration spent in new arm of Y maze. (B) The number of entries to new arm of the Y maze. Columns, mean; bars, SEM.

