## Supplemental Figures for "The BMP antagonist *Gremlin1* contributes to the development of cortical excitatory neurons, motor balance and fear responses"

### Ichinose_Supplemental_Fig_S1

A

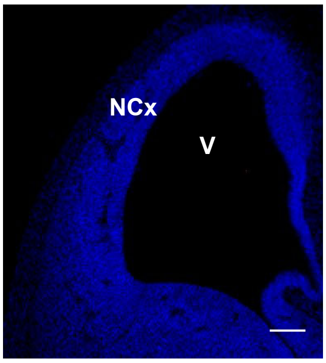


B

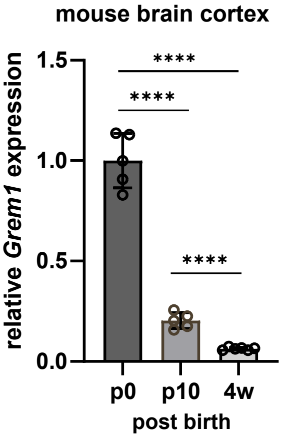

### Ichinose_Supplemental_Fig_S2

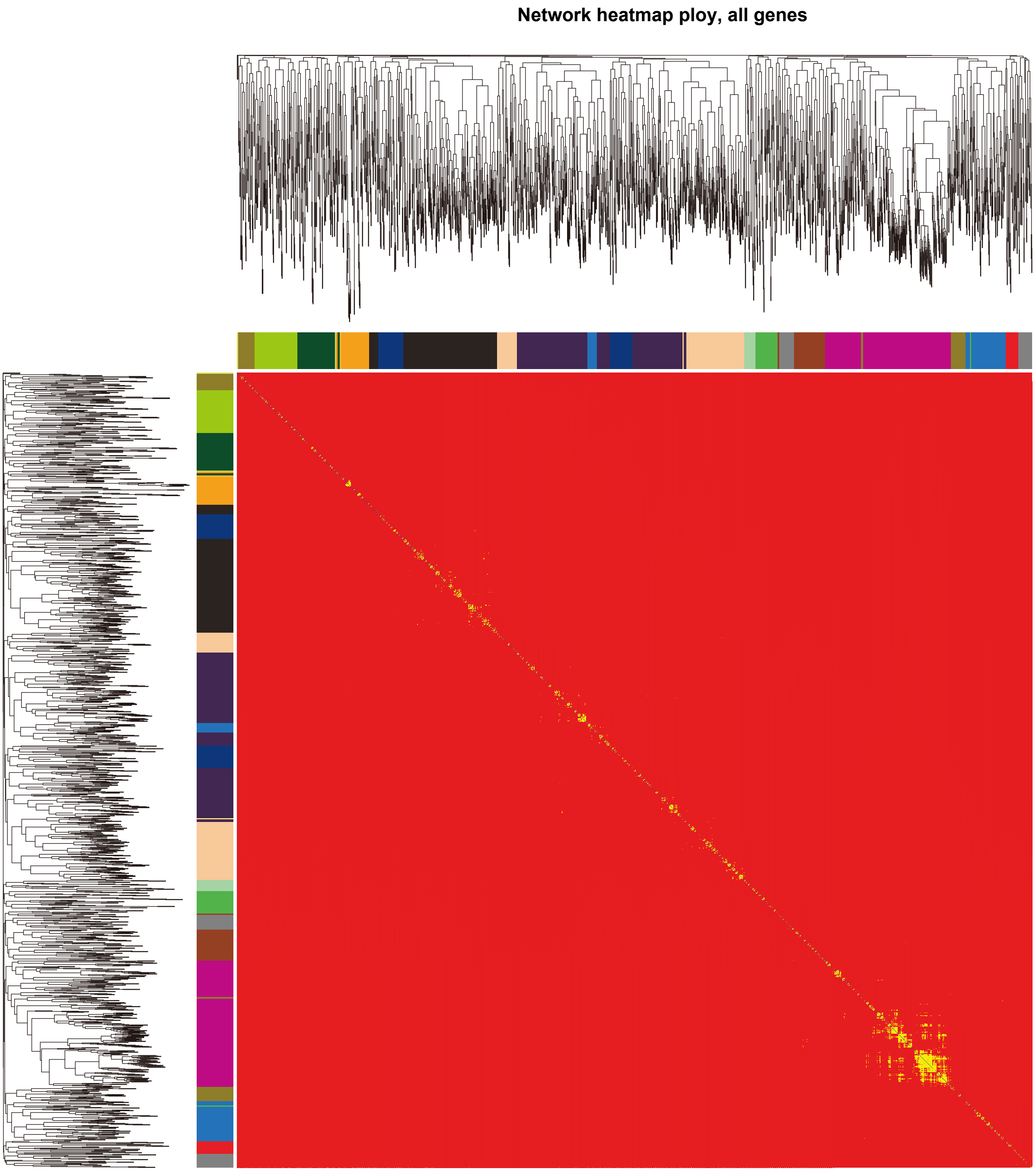

**Ichinose_Supplemental_Fig_S3**

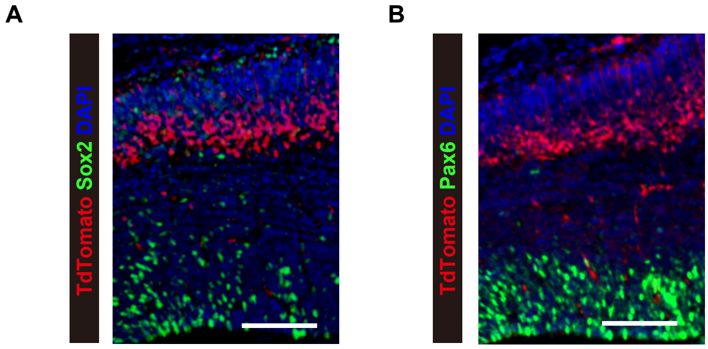

### Ichinose_Supplemental_Fig_S4

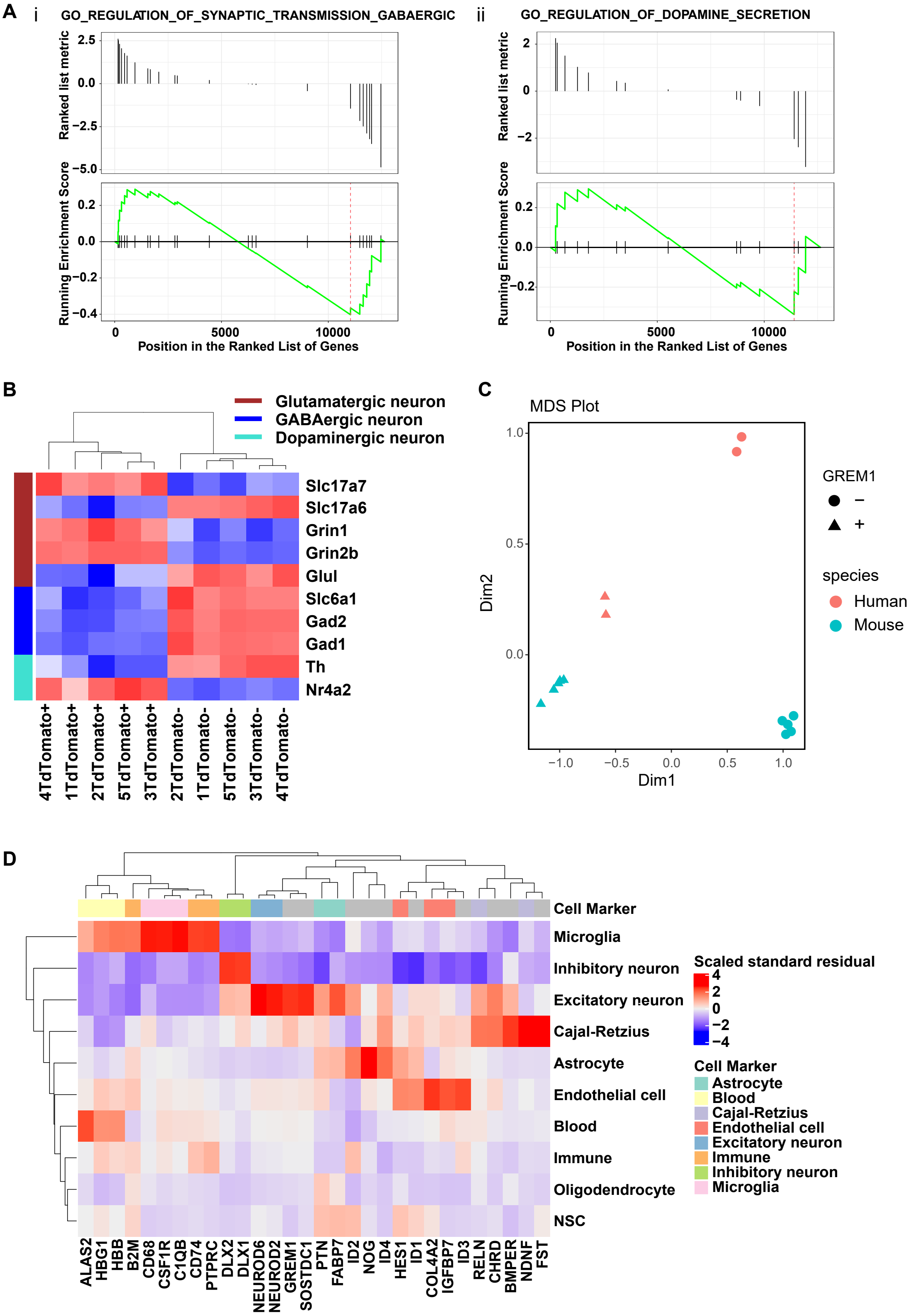

Ichinose_Supplemental_Fig_S5

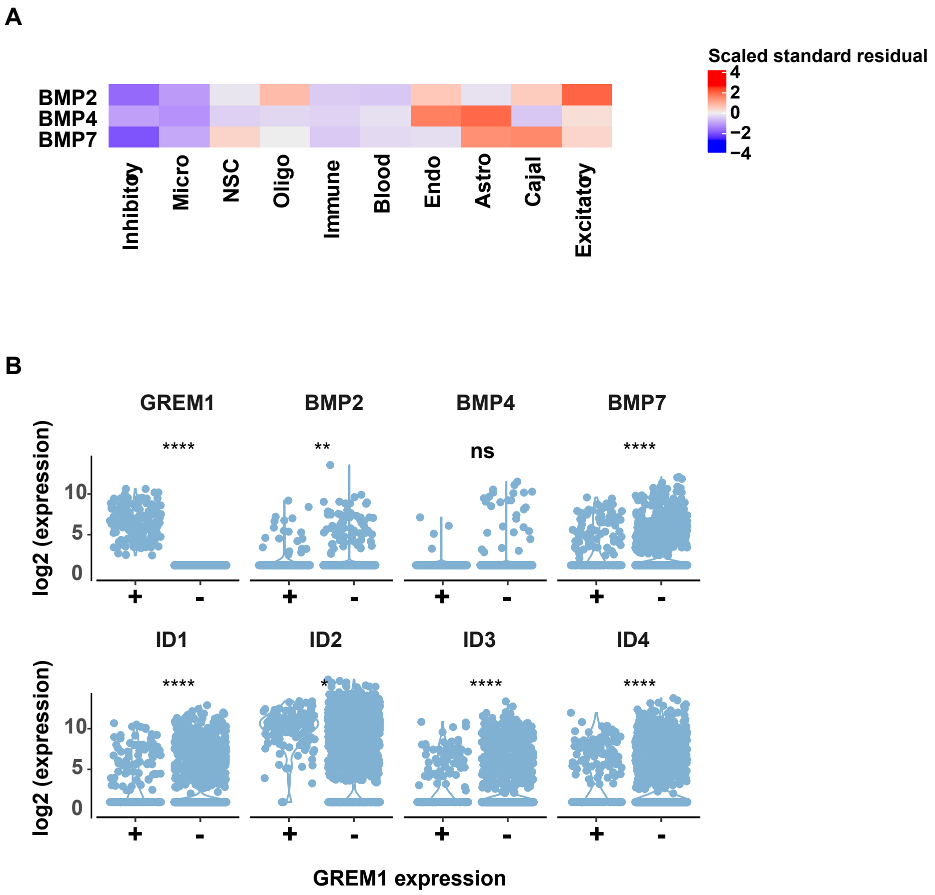

### Ichinose_Supplemental_Fig_S6

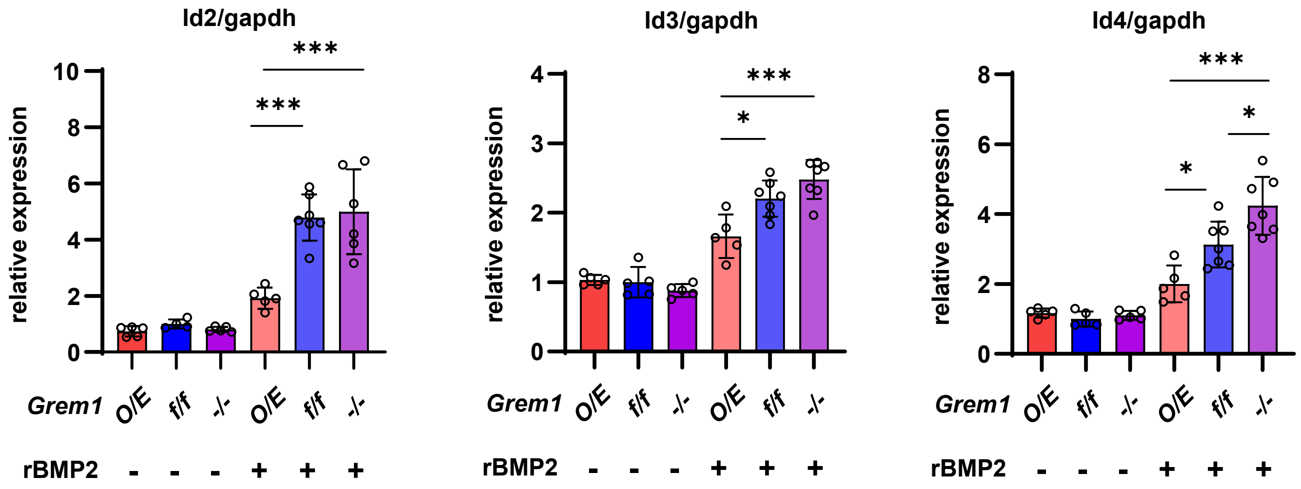

### Ichinose_Supplemental_Fig_S7

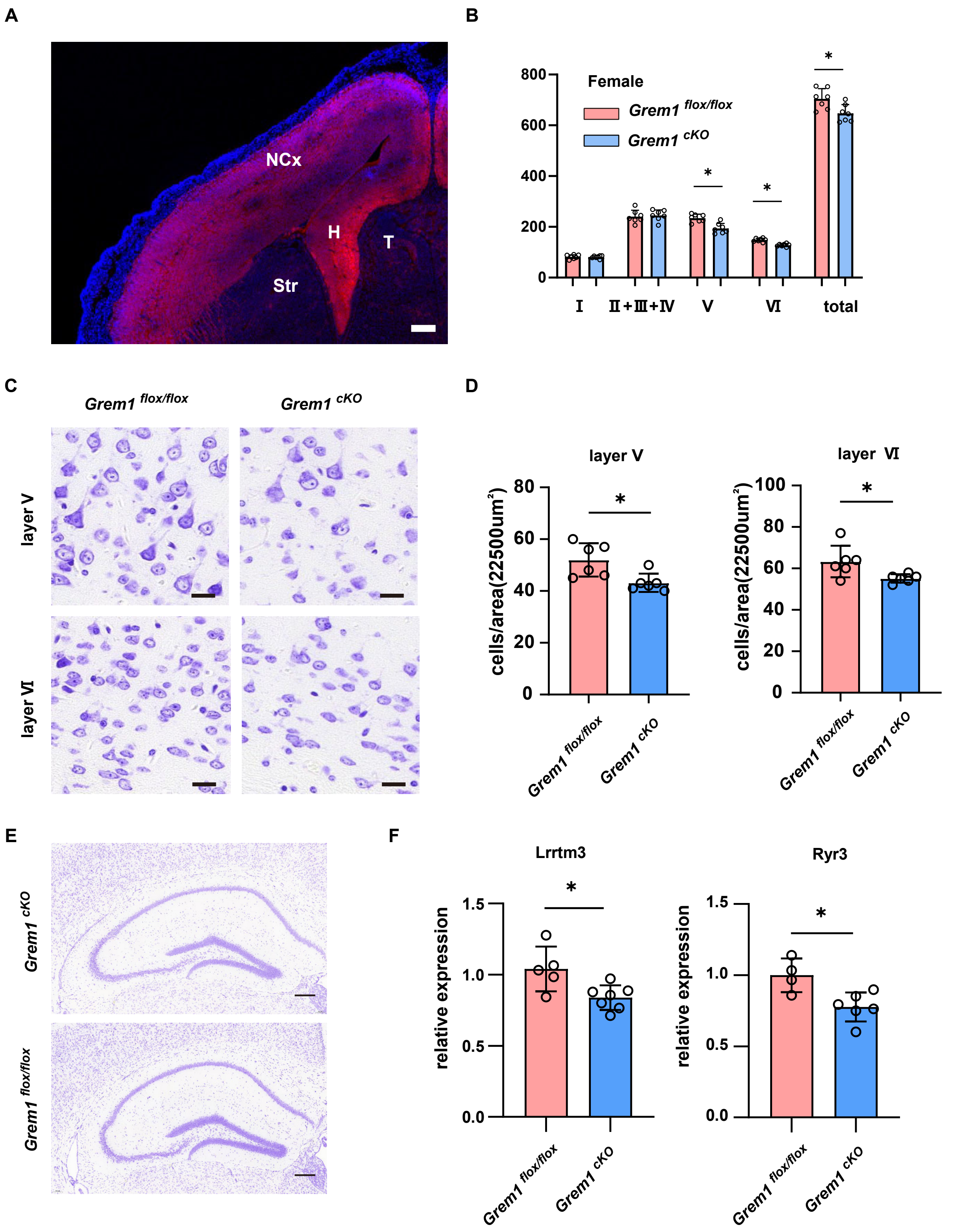

### Ichinose_Supplemental_Fig_S8

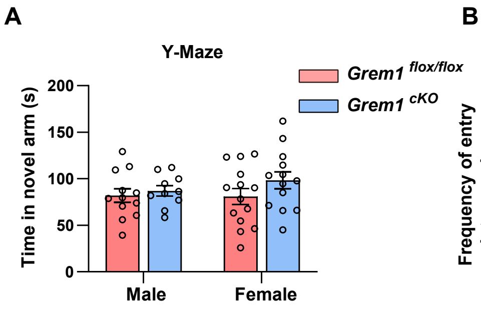


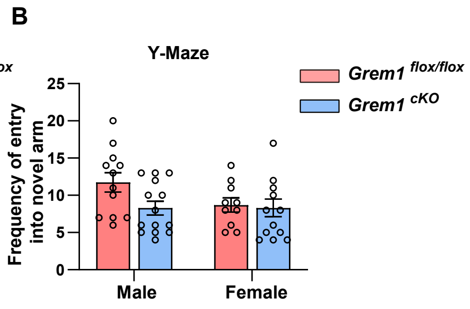
